# The contribution of neutrophils to bacteriophage clearance and pharmacokinetics in vivo

**DOI:** 10.1101/2024.01.25.577154

**Authors:** Arne Echterhof, Tejas Dharmaraj, Robert McBride, Joel Berry, Max Hopkins, Hemaa Selvakumar, Lynn Miesel, Ju-Hsin Chia, Kun-Yuan Lin, Chien-Chang Shen, Yu-Ling Lee, Yu-Chuan Yeh, Wei Ting Liao, Gina Suh, Francis G. Blankenberg, Adam R. Frymoyer, Paul L. Bollyky

## Abstract

With the increasing prevalence of antimicrobial-resistant bacterial infections, there is great interest in using lytic bacteriophages (phages) to treat such infections. However, the factors that govern bacteriophage pharmacokinetics *in vivo* remain poorly understood. Here, we have examined the contribution of neutrophils, the most abundant phagocytes in the body, to the pharmacokinetics of intravenously administered bacteriophage in uninfected mice. A single dose of LPS-5, an antipseudomonal bacteriophage recently used in human clinical trials, was administered intravenously to both wild-type BALB/c and neutropenic ICR mice. Phage concentrations were assessed in peripheral blood and spleen at 0.5, 1, 2, 4, 8, 12, and 24 hours after administration by plaque assay and qPCR. We observed that the phage clearance is only minimally affected by neutropenia. Indeed, the half-life of phages in blood in BALB/c and ICR mice is 3.45 and 3.66 hours, respectively. These data suggest that neutrophil-mediated phagocytosis is not a major determinant of phage clearance. Conversely, we observed a substantial discrepancy in circulating phage levels over time when measured by qPCR versus plaque assay, suggesting that substantial functional inactivation of circulating phages occurs over time. These data indicate that circulating factors, but not neutrophils, inactivate intravenously administered phages.

## Introduction

The global crisis of bacterial antimicrobial resistance (AMR) represents a profound challenge to human health. Once-treatable infections are now responsible for extensive morbidity and mortality and have the potential to evolve into global health problems of pandemic dimensions^1^.

*Pseudomonas aeruginosa* (*Pa*) is one of the most problematic pathogens for which new treatment options are needed^2-4^. Multidrug-resistant strains of *Pa* are prevalent in environmental and clinical settings due to acquired resistance and intrinsic mechanisms of resistance^5,6^. There is a need for innovative treatments to treat AMR *Pa* and other pathogens. Lytic bacteriophages (phages), viruses that kill bacteria, are an exciting treatment option for AMR bacterial infections^7-10^. Multiple studies have demonstrated the efficacy of phage therapy against MDR *Pa*^*11-14*^, either alone or in conjunction with conventional antibiotics^15^. Phage therapy is saving lives; however, success has been inconsistent^16-18^. This has limited the therapeutic and commercial prospects of this approach.

Critical to the successful development of phages as a therapeutic treatment will be a deep understanding of its pharmacokinetics (PK)^19^. While there is, in fact, a large body of literature on phage therapy, it is heterogenous, difficult to access, as much of it is in the non-English-language literature, and often pre-dates the current era of phage therapy. Fortunately, this literature has been summarized in an excellent recent review^20^.

From this literature, it is clear that phage PK depends on the route of administration. Following intravenous (IV) administration, the half-life (*t*_1/2_) of phages in the blood of mice has been reported between 2.2 h and 4.5 h in different animal models^21,22^. Most phages are cleared by the liver and spleen within minutes to hours^23-25^. However, systemic administration of phages leads to their accumulation in tissues. The removal of phages from circulation and their clearance decreases the number of infective phage particles and affects therapeutic efficacy^26^. However, the factors responsible for phage clearance are unclear. This is important to understand in order to better optimize phage delivery and treatment protocols.

The immune system has been implicated in phage clearance with phagocytosis playing a prominent role in the clearance of many viruses and foreign molecules. Large viral particles exit the circulation and enter the liver and splenic sinusoids, where a high density of phagocytes are present^26,27^. Nanosized virions are destroyed in the reticuloendothelial system (RES) of the liver and spleen, which leads to rapid removal of phages from circulation. While in the reticuloendothelial system, phages may also exert pharmacodynamic effects by influencing the inflammatory properties of mononuclear cells^22,28,29^.

Neutrophils are the most abundant phagocytes in the body and can be found in the liver, spleen, and in peripheral tissues. Neutrophils have been reported to engulf viruses^36–38^. However, the contributions of neutrophils and phagocytosis to phage clearance remains unclear.

In this study, we evaluated the pharmacokinetics of intravenously administered *Pseudomonas aeruginosa* phage LPS-5 in a wild-type and neutropenic mouse model.

## Materials and Methods

### Chemicals and reagents

The following materials were used in these studies: 0.9% NaCl (Sin-Tong, Taiwan), 2X Power SYBR Green PCR Master, Bacto™ Agar (214040, BD, USA), Disodium phosphate (Na_2_HPO_4_, 71640, Sigma, USA), LB Broth (Lennox) (240230, BD, USA), Magnesium sulfate (MgSO_4_, M2643, Sigma, USA), Nutrient agar (NA) plates (CMP0101312, CMP, Taiwan), Nutrient broth (DIFCO, USA), Phosphate buffered saline (PBS) tablet, pH 7.4 (P4417, Sigma, USA), Potassium phosphate monobasic (KH_2_PO_4_, P5379, Sigma, USA), Sodium chloride (NaCl, S7653, Sigma, USA), TANBead nucleic acid extraction kit (M665A46, OptiPure Viral Auto plate, Taiwan), Tryptic soy broth (TSB) (211825, BD, USA), and Water for injection (Tai-Yu, Taiwan).

### Animals

Specific pathogen-free immunocompetent female BALB/c mice, 7-8 weeks of age, were used for the pharmacokinetics study. Animals were housed for 3 days in quarantine prior to the pharmacokinetics study. Specific pathogen-free neutropenic female Institute of Cancer Research Hsd:ICR (ICR) mice, weighing 22 ± 2 g were used. Animals were immunosuppressed by two intraperitoneal injections of cyclophosphamide, the first at 150 mg/kg 4 days before infection (Day –4) and the second at 100 mg/kg 1 day before infection (Day –1), prior to infection on Day 0 per the industry-standard methods^30^. Pharmacology Discovery Services, PDS, determined that this cyclophosphamide treatment schedule resulted in neutropenia (<100 neutrophils per µL) until Day 2 after infection.

### Phage suspension

Phages were propagated using techniques that are well-established at Felix Biotechnology (South San Francisco, CA). Briefly, we infected planktonic cultures of bacteria with phage until clearing was observed relative to a non-infected control culture, removed gross bacterial debris by centrifugation (8000 x *g*, 20 min, 4 °C), and filtered the supernatant through a 0.22 um PES membrane (Corning, Corning, New York, Product #4311188). The supernatant was treated with 5 U/mL Benzonase nuclease (Sigma-Aldrich, Saint Louis, MO, Catalog #E8263) overnight at 37 °C to digest free DNA. Phage was precipitated by the addition of 0.5 M NaCl + 4% w/v polyethylene glycol 8000 (Sigma-Aldrich, Saint Louis, MO, Catalog #PHR2894) overnight at 4 °C. Precipitated phage was then pelleted by centrifugation (14,000 x *g*, 20 min, 4 °C), washed in 30 mL Tris-EDTA buffer (10 mM Tris-HCl, 1 mM EDTA, pH 8.0), and re-pelleted by centrifugation (14,000 x *g*, 20 min, 4 °C). The phage pellet was then resuspended in 3 mL of the buffer appropriate to that phage/experiment and dialyzed against 4 L of the same buffer three times through a 10 kDa dialysis membrane to remove residual salts and PEG. Felix Biotechnology supplied the LPS-5.eng bacteriophage stock solution at 1.0 x 10^11^ PFU/mL and the heat-inactivated phage solution.

PDS then stored the phages at 4 °C upon arrival at PDS. Dosing solutions for injection were prepared by diluting the stock solution in filtered phage buffer (KH_2_PO_4_ 144 mg/L, Na_2_HPO_4_ 421.62 mg/L, NaCl 9000 mg/L, MgSO_4_ 1203.6 mg/L) within 1 h before administration on each dosing day. The phage dose solutions were titered twice on two separate test occasions with the plaque method.

### Treatment

LPS-5 phage was administered intravenously via tail vein injection at a single dose (0.1 mL/mouse with a concentration of 1 x 10^11^ PFU/mL) to BALB/c mice or ICR mice. Animals were observed at 5-15 min after IV dosing to detect acute toxicity, which would have been recorded. No toxicity was observed. Animals were then sacrificed after 0.25 h, 1 h, 2 h, 4 h, 8 h, 12 h and 24 h. For every sampling time point, five mice were sacrificed. Phage concentration was measured in whole blood, liver and spleen homogenates using a plaque assay and qPCR.

### Blood and Tissue Collection

Animals were euthanized via CO_2_ euthanasia at the time points noted for blood and tissue collection. Blood samples were collected by cardiac puncture under CO_2_ euthanasia using lithium heparin blood collection tubes. The blood samples were stored on ice and divided into two vials, one for PFU enumeration and one for quantitative PCR (qPCR).

Tissues were aseptically recovered, blotted dry, weighed, then combined with 1 mL PBS followed by homogenization with a Polytron homogenizer (10,000 x *rpm* for 15-30 seconds). The homogenized samples were centrifuged and the supernatant was transferred to two vials, one for PFU enumeration and one for quantitative PCR (qPCR). All samples were stored on ice during processing which was completed within one hour.

### Plaque assays

Plaque assays were used to quantify the number of infectious phage particles per mL. Serial 10-fold dilutions of plasma and clarified homogenates were prepared in phage dilution buffer. The assay was conducted on rectangular OneWell bottom agar plates (128 x 86 mm), containing LB agar (1.5%) with 10 mM MgSO_4_. Seeded top agar, ∼10 mL, was spread evenly over the upper surface of the bottom agar plate and was prepared by combining 300 µL of overnight culture of *P. aeruginosa* (PAO1) with 10 mL molten top agar medium (LB broth medium with 10 mM MgSO_4_ and 0.75% agar). The overnight culture of strain *P. aeruginosa* (PAO1) was grown in LB with 10 mM MgSO_4_. After solidification, 5 µL of each sample dilution was pipetted onto the surface of the top agar, then left to dry. The plate was incubated at 37 °C for 18 h then plaques were counted and PFU/mL values were calculated. Plaques were counted on the spots with 1 to 50 distinguishable plaques to calculate PFU/mL values. The number of plaques per dilution was tabulated and the PFU/tissue or PFU/mL blood was calculated and reported.

### Transmission electron microscopy (TEM)

TEM imaging was done as previously reported^31^. In brief, the size and morphology of phages were examined with transmission electron microscopy (TEM) using a JEOL JEM1400 (JEOL USA Inc., Peabody, MA) at 80 kV. 5 µL of diluted phage solution was dropped onto carbon-coated copper grids (FCF-200-Cu, Electron Microscopy Sciences, Hatfield, PA). After 3 minutes, the grid was dipped into a ddH_2_O droplet and then 1% uranyl acetate for staining was dropped at the sample and was allowed to dry for 15 minutes before performing microscopy.

### Quantitative PCR (qPCR)

Sample Preparation: Nucleic acid was extracted from the collected blood samples, and clarified tissue homogenates using the TANBead Nucleic Acid Extraction kit with the Maelstrom 4810 fully automated DNA/RNA extraction system following the manufacturer’s instructions

Quantitative PCR: qPCR was conducted with 2X Power SYBR Green PCR Master Mix, Thermo Fisher, USA using the standard assay components of the kit. The qPCR reactions were performed on a ViiA™ 7 Real-Time PCR System (Applied Biosystems) using a 96-well format. Assay wells were sealed with optical films and then briefly centrifuged to consolidate the droplets from the sides of wells. Thermal cycling included an initial 10-min heat activation of the polymerase, followed by 40 repeated cycles of DNA denaturation, primer annealing, and target elongation. Two primer pairs with sequences supplied by Felix Biotechnology were used for the qPCR. The MCP_set2 primer set targets nucleotides 5223 to 5367 of the phage and yields a 145 bp product with a melting temperature of 81.51 °C. The Barcode primer set targets a unique barcode in the genetically modified phage at nucleotides 70423 to 70750 and yields a 328 bp product with a melting temperature of 85.34 °C. MCP_set2_F: CGCAACTGGGCTAACACCGC and R: GGTTGGTCAGGTCGAAGCC Barcode_F_oFB123: GTGCAGGAGGAATCGGGCC and R_oFB121: GGGTCACTTTCACTTGCACGC.

Condition optimization: qPCR condition optimization by analysis of melting temperatures was conducted to inspect for the specificity of the amplified product. To assess specificity, qPCR was performed with primer pairs for each phage using nucleic acid prepared from infected *P. aeruginosa* mouse blood. The qPCR signal and the threshold cycle value (CT) was measured in triplicate in each of these samples and the signal-to-background values were determined. Tests were conducted to confirm that the primers would only yield amplicons from template samples that contained phage DNA. Template samples containing mouse blood or bacteria, but lacking phage, were tested with the expectation that they would not yield an amplification signal. A dilution series of the phage were tested to define the linear range of the qPCR assay for each phage, the lower limit of detection and quantification (LLOD and LLOQ), the upper limit of quantification (if measurable, ULOQ), and the efficiency of amplification, The experiment assessed the linearity of the qPCR signal from the phage over ten concentrations: 0, 10, 25, 50, 10^2^, 10^3^, 10^4^, 10^5^, 10^6^, 10^7^, and 10^8^ PFU. Blood samples and homogenized tissue samples were spiked with phage dilutions at these ten concentrations to mimic samples from treated animals. Mouse biosamples without spiked phage were included as negative controls. DNA was extracted and measured as described above. The qPCR analysis of each sample was conducted in triplicate to assess intra-assay variability. *qPCR standard curve*

Calibration curve samples were generated by spiking homogenized tissues or blood samples with phage dilutions of defined concentrations in an 8-point dilution series, ranging from 10^1^ to 10^8^ PFU/mL. Triplicate qPCR measurements of each phage were conducted for each sample. For calibration curve samples, the CT values were graphed as a function of the nominal phage concentration (data not shown). The concentration of each phage in the tissue sample of unknown concentration was determined by comparing the measured CT values against the calibration curve from the same biological matrix, per the formula below.

Each qPCR condition included four QC samples that are prepared according to the same extraction procedure and the same test occasion as the test samples. Clarified tissue or blood homogenates, were spiked at four concentrations: the LLOQ, low, mid, and high concentrations of phage, three replicate samples. Controls included no phage vehicle samples. The no-phage blanks were evaluated for the presence of signal artifacts due to contamination.

The CT values of the dilution samples were graphed as a function of the logarithm of the nominal phage concentration (PFU/mL). The amplification efficiency, E, was determined from the slope per the equation below. Regression analysis was conducted to determine the r values to assess linearity. Concentrations of phage (PFU/mL) in each sample were back-calculated from the calibration standard curve, per the equation below. The calculated concentrations were compared to the nominal concentrations to calculate the assay precision.

Efficiency = 10^-1/slope^ - 1

DNA phage equivalence (PFU/mL or PFU/tissue) = 10^(Ct value – Yintercept)/slope^

Accuracy: measured/nominal*100%

Precision: SD/Mean*100%

Acceptance criteria: The dilution series is to yield an r value of 0.98 or higher over seven or more concentrations. The precision is to be 85-115% for 70% of the calibration curve samples. The amplification efficiency is to be 85-105%. The linear range should be between 10^3^ to 10^8^ PFU/mL or PFU/tissue of samples.

The concentration of each phage titer was determined in the blood and tissue samples. The concentration-time profile of phage measured with qPCR was characterized as described above for the plaque assay. The outcomes of the qPCR and the phage titration assays were compared.

### PK-data analysis

Plots of phage concentration in blood, spleen, and liver tissues over time were generated with GraphPad Prism software. Non-compartmental analysis (NCA) was conducted with Phoenix WinNonLin to estimate pharmacokinetic parameters. C_max_ and C_24h_ were taken directly from the data. The terminal elimination rate (λ_z_) was determined by linear regression analysis of the terminal portion of the log plasma concentration–time curve. The terminal half-life (*t*_1/2_) was calculated as ln 2/λ_z_. AUC_last_ was found using the trapezoidal method. Summation of AUC_last_ plus the concentration at the last measured point divided by λ_z_ yielded AUC_Inf_. CL was calculated as dose/AUC_Inf_. The mean residence time (MRT) was calculated as the ratio of the area under the moment curve (AUMC_Inf_ divided by AUC_Inf_. Volume of distribution based on the terminal phase (V_z_) was calculated as Dose/(λ_z_ x AUC_inf_).

## Results

### Calibration studies to evaluate our ability to assess phage concentrations

As the phage in this study, we used LPS-5, a lytic phage recently included in a human clinical trial of phage therapy, the CYstic Fibrosis bacterioPHage Study at Yale (CYPHY) (ClinicalTrials.gov Identifier: NCT04684641). LPS-5 is a lytic phage that infects *Pa*. It is a member of the *Pakpunavirus* family and uses pseudomonal lipopolysaccharide (LPS) for viral entry^14^. A representative transmission electron microscopy image of LPS-5 and representative lytic plaque morphology on *Pseudomonas aeruginosa* PAO1 are shown in **Fig. 1**.

**Figure 1:**
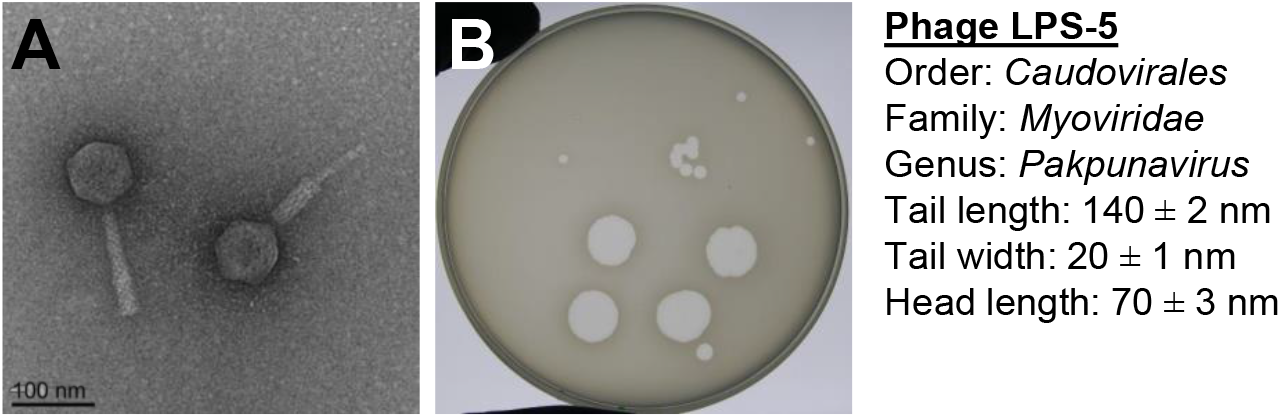
LPS5, the phage used in these studies. **A**. A representative transmission electron microscopy image of LPS-5 is shown at 80,000x magnification. **B**. Lytic plaque morphology for LPS-5 on *Pseudomonas aeruginosa* PAO1.

We first sought to assess the linearity, accuracy, and precision of our methods to quantify phages in serum and tissues collected from conventional BALB/c mice. To assess this, we performed titration studies to determine the accuracy and precision of phage measurement over a broad phage range of concentrations. For these studies, previously collected peripheral blood and excised spleen tissue were spiked with phage in eight concentrations (10^8^, 10^7^, 10^6^, 10^5^, 10^4^, and 10^3^ PFU/sample). For spleen tissues, the phages were spiked in, and these were homogenized together at 10,000 x *rpm*. For blood, the phages were added into whole blood and mixed. As readouts for these studies, we used plaque assays and qPCR studies, as detailed in the Material & Methods section. For these studies. phages with established, nominal concentrations were added to PBS (saline), blood, or spleen tissue collected from BALB/c mice (A-F) or ICR mice (G-L). These were then analyzed using plaque assays (A-C, G-I) or qPCR (D-F, J-L) **(Fig. 2)**.

**Figure 2:**
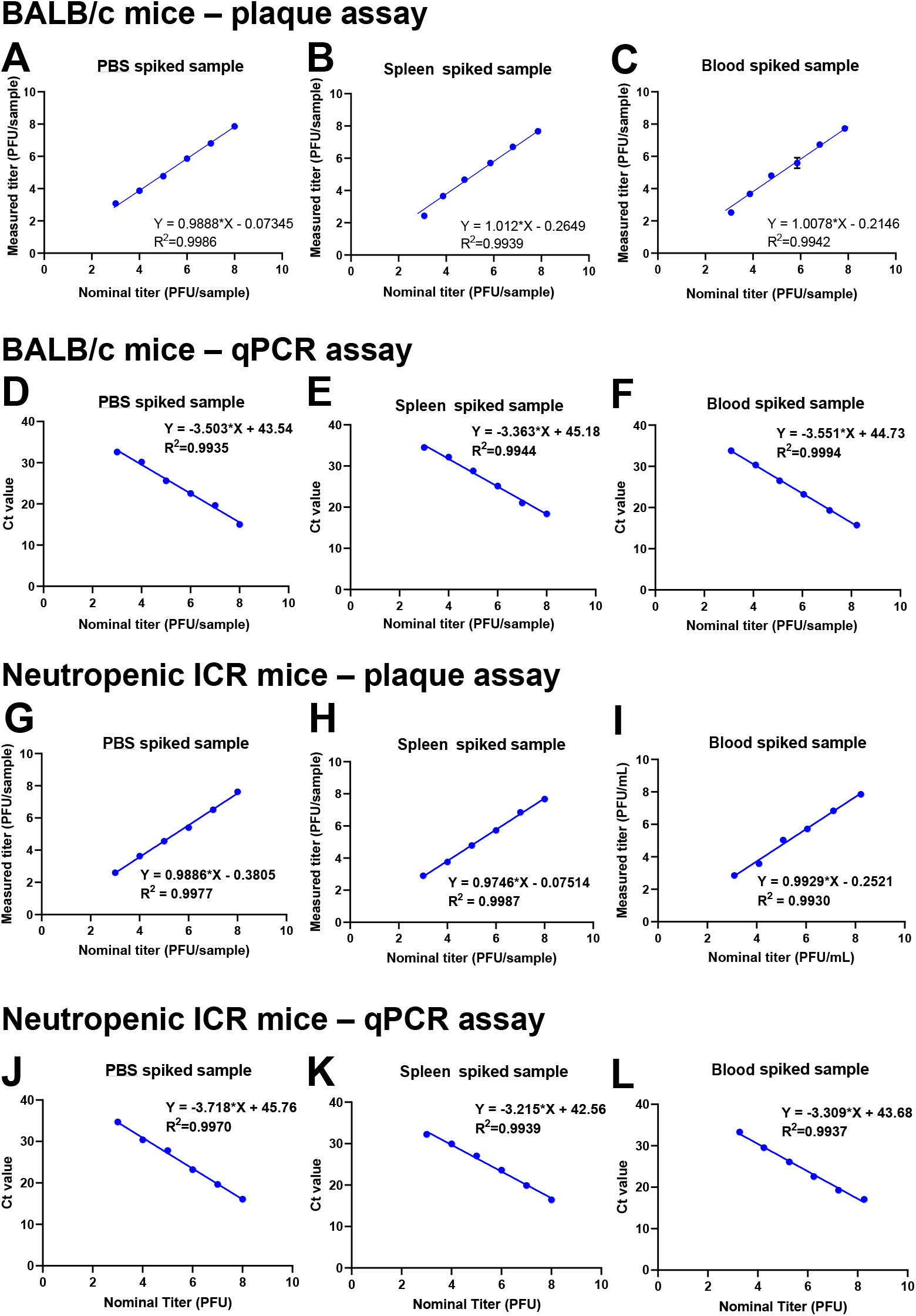
Calibration curves for phage titrations in blood and spleen. To assess our ability to accurately quantify phages in blood and tissue samples, we performed titration studies using plaque assays and qPCR assays as our readouts. For these studies phages with established, nominal concentrations were added to PBS (saline), blood, or spleen tissue collected from BalbC mice (**A-F**) or ICR mice (**G-L**). These were then analyzed using plaque assays (**A-C, G-I**) or qPCR (**D-F, J-L**).

For the qPCR studies, the accuracy in BALB/c samples was determined to be 100% (99-121) (spleen) and 101% (94-104) (blood). The uncertainty was determined to be 0.7% (0.2-2.0) (spleen) and 0.4% (0.2-3.0) (blood). The accuracy in ICR samples was determined to be between 87% (81-92) (spleen) and 97% (93-99) (blood). The uncertainty was determined to 0.3% (0.2-1.5) (spleen) and 0.4% (0.2-3.0) (blood). The limit of detection (LOD) was determined to be 200 PFU/sample. The lower limit of qualification (LLOQ) was determined to be 1000 PFU/sample. These are indications of excellent linearity and reproducibility using these approaches.

### The PK of phage LPS-5 in conventional BALB/c mice

Having established our ability to assess phage levels in these samples, we next asked whether we could also assess the pharmacokinetics of phages in vivo. For these studies, we used a well-established mouse commonly used in pharmacology studies, the BALB/c mouse.

Using these animals, we then examined phage pharmacokinetics using the protocol outlined in **Fig. 3**. In these studies, LPS-5 Phage was IV administered at a single dose (0.1 mL/mouse with a concentration of 1 x 10^11^ PFU/mL) to BALB/c mice. Animals were then sacrificed after 0.25 h, 1 h, 2h, 4 h, 8 h, 12 h and 24 h. For every sampling time point, five mice were sacrificed. Overall, 45 immunocompetent mice were used. Phage concentration was measured in whole blood and spleen homogenates using a plaque assay and qPCR. The concentration over time was graphed and non-compartmental analysis (NCA) was conducted with Phoenix WinNonLin to estimate pharmacokinetic parameters.

**Figure 3:**
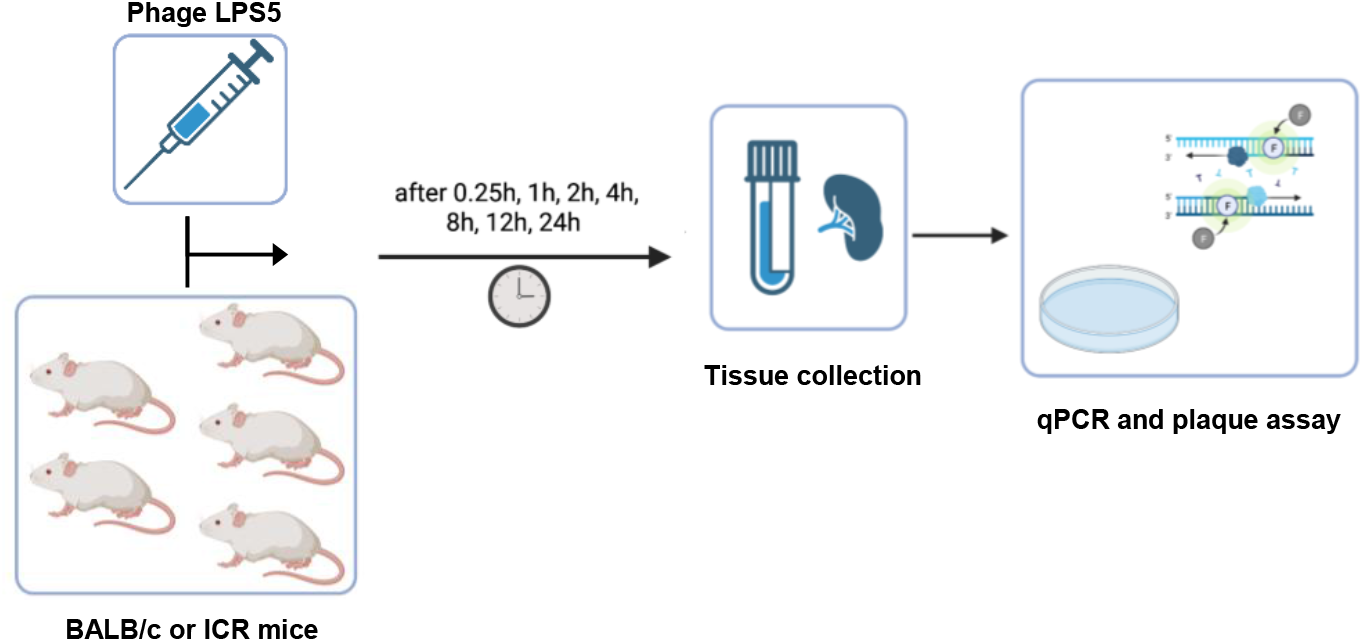
Schematic of biodistribution and pharmacokinetics experiment. In these studies, Phage LPS5 was administered IV as a single dose (0.1 mL/mouse at 1×10_11_ PFU/mL) to BALB/c mice or ICR mice. Animals were sacrificed after 0.25, 1, 2, 4, 8, 12 or 24. At every sampling time point, five mice were sacrificed, and the phage concentration was measured in whole blood and spleen homogenates by qPCR and plaque assay. Non-compartmental analysis (NCA) was conducted with Phoenix WinNonLin to estimate pharmacokinetic parameters.

The concentration-time profile as measured by plaque assay of active LPS-5 after IV administration in conventional BALB/c mice is shown **Fig. 4A**. The C_max_ of active phages in blood in BALB/c mice was 1.97 x 10^6^ PFU/mL. Circulating active phages then demonstrated first order elimination with a terminal *t*_1/2_ of 3.45 h. By 24 h, very low concentrations of active phage remained in the blood (C_24_ = 740 PFU/mL). In contrast, high concentrations of active phages in spleen tissue were rapidly achieved (C_max_ = 5.21 x 10^8^ PFU/mL) and remained high over the 24 hours period of measurement (C_24_ = 6.46 x 10^7^ PFU/mL) **(Fig. 4B)**. Other pharmacokinetic parameters are listed in **Table 1**.

**Table 1:** Pharmacokinetic parameters for BALB/c and neutropenic ICR mice in blood and spleen based on qPCR ad plaque assay. The pharmacokinetic parameters were obtained from non-compartmental analysis (NCA) using Phoenix WinNonlin software. The following values were calculated: elimination half-life t_1/2_, Initial concentration 0.25h after dose C_0_, concentration after 24h C_24_, Area under the curve from the time of dosing to the time of the last measurable concentration AUC_last_, AUC from the time of dosing extrapolated to infinity AUC_Inf_, Mean residence time MRT, Volume of distribution at steady state V_ss_, and clearance CL

| Parameter | BALB/c mice;<br>Blood |  | ICR mice;<br>Blood |  | BALB/c mice;<br>Spleen |  | ICR mice;<br>Spleen |  |
| --- | --- | --- | --- | --- | --- | --- | --- | --- |
|  | qPCR | Plaque assay | qPCR | Plaque assay | qPCR | Plaque assay | qPCR | Plaque assay |
| $t_{1/2}$ (h) | 16.26 | 3.45 | 9.15 | 3.66 | 6.11 | 11.54 | 8.21 | 5.18 |
| $C_{MAX}$ (PFU/mL) | $5.58 \times 10^6$ | $1.97 \times 10^6$ | $3.99 \times 10^6$ | $2.19 \times 10^7$ | $4.37 \times 10^8$ | $5.71 \times 10^8$ | $3.41 \times 10^8$ | $3.89 \times 10^8$ |
| $C_{24}$ (PFU/mL) | $2.73 \times 10^4$ | 740 | 3610 | 280 | $3.99 \times 10^7$ | $6.46 \times 10^7$ | $3.45 \times 10^7$ | $5.31 \times 10^6$ |
| Cl (1/h) | $2.41 \times 10^9$ | 6730 | 282 | 86.3 | 2.47 | 1.81 | 3.63 | 8.35 |
| $V_{ss}$ (1/mL) | $2.49 \times 10^4$ | $1.37 \times 10^4$ | 411 | 140 | 25.6 | 27.0 | 44.8 | 55.6 |
| $AUC_{last}$ (PFU h/mL) | $3.50 \times 10^6$ | $1.48 \times 10^6$ | $3.54 \times 10^7$ | $1.16 \times 10^8$ | $4.04 \times 10^9$ | $5.53 \times 10^9$ | $2.75 \times 10^9$ | $1.20 \times 10^9$ |
| $AUC_{inf}$ (PFU h/mL) | $4.14 \times 10^6$ | $1.49 \times 10^6$ | $3.54 \times 10^{-3}$ | $1.16 \times 10^{-2}$ | 0.404 | 0.553 | 0.275 | 0.120 |
| $AUC_{extr}$ (%) | 15.48 | 0.25 | 0.13 | 0 | 8.7 | 19.42 | 14.84 | 3.31 |
| MRT (h) | 10.33 | 2.03 | 1.46 | 1.62 | 10.34 | 14.95 | 12.32 | 6.66 |
| $r^2$ | 0.9998 | 0.986 | 0.7287 | 0.925 | 0.971 | 0.6815 | 0.9792 | 0.9591 |
|  | qPCR |  | qPCR |  | Plaque assay |  | Plaque assay |  |
| Spleen/blood $C_{24}$ | $1.46 \times 10^3$ | | $9.56 \times 10^3$ | | $8.72 \times 10^4$ | | $1.90 \times 10^4$ | |
| Spleen/blood $AUC_{last}$ | $1.05 \times 10^3$ | | 66.25 | | $3.00 \times 10^3$ | | 10 | |

**Figure 4.**
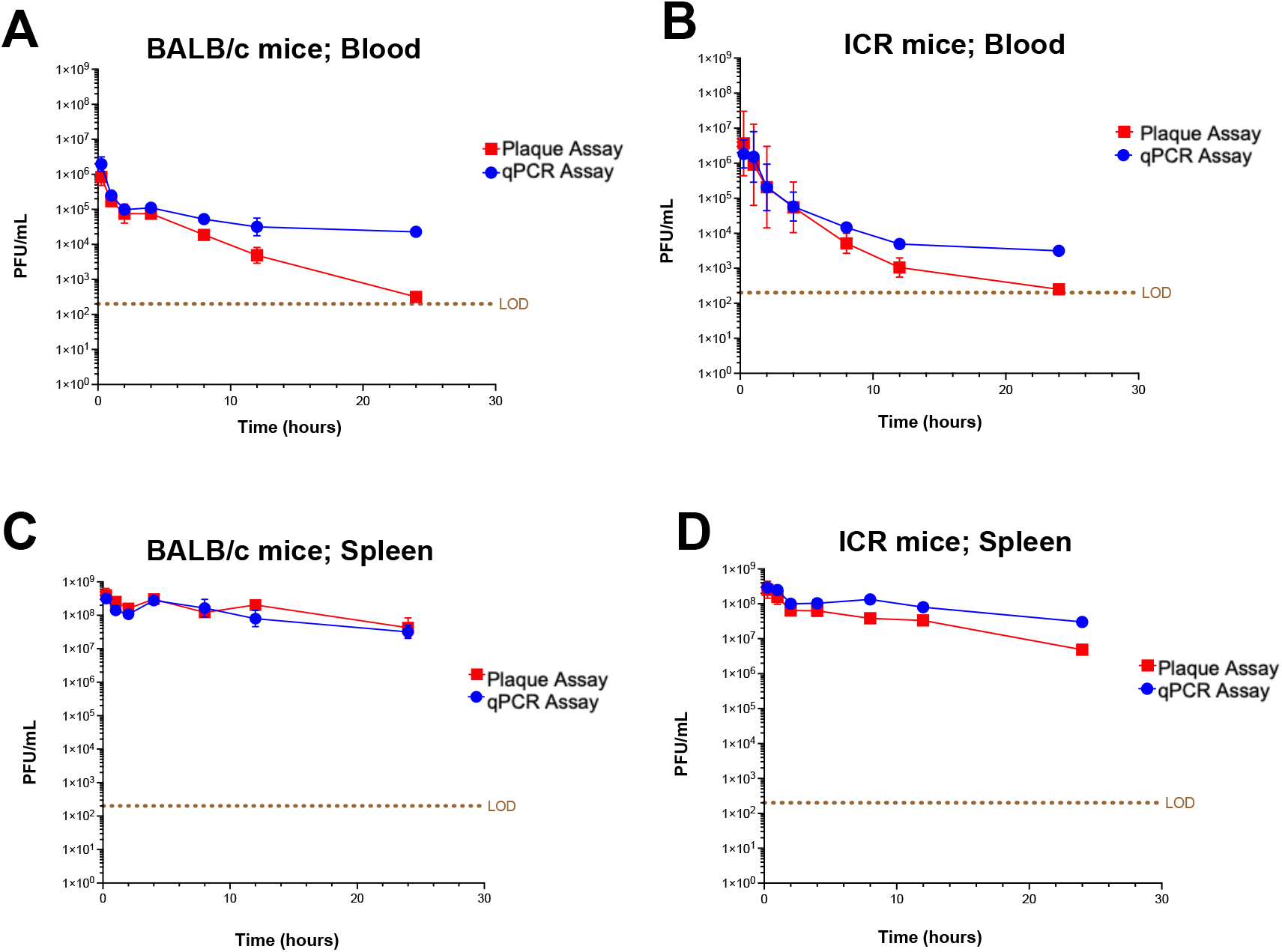
Pharmacokinetics and biodistribution of LPS5 bacteriophage in blood and spleen tissue over time in BalbC (wild type) and ICR (neutropenic) mice. Mice were injected with 0.1 mL of an 1×10_10_ PFU/mL LPS5 phage suspension. At each sample time point over 24 h, five mice were sacrificed at the 0.25, 1, 2, 4, 8, 12, and 24 h. qPCR and plaque assays were performed on blood and spleen tissue homogenates. 45 immunocompetent mice and 45 neutropenic mice were used (for experimental design, see Material and Methods). Mean +/- SD is plotted over time. **(A, C)** Blood pharmacokinetics of LPS5 in immunocompetent BALB/c (**A**) and neutropenic ICR (**C**) mice. **(B, D)** Spleen biodistribution of LPS5 in immunocompetent BALB/c (**B**) and neutropenic ICR (**D**) mice. LOD = limit of detection.

Together, these data suggest that bacteriophages leave the circulation rapidly after administration and accumulate in reticuloendothelial system compartments like the spleen.

### The PK of phage LPS-5 in neutrophil-deficient mice

We then asked how neutropenia impacted phage PK and biodistribution. For these studies, we used the ICR neutropenic mouse model where ICR mice are treated with cyclophosphamide to render them neutropenic, as detailed in the methods section. These animals are often used to study pharmacokinetics^39,40^. Here again, in these studies, LPS-5 phage was IV administered at a single dose (0.1 mL/mouse with a concentration of 1 x 10^11^ PFU/mL) to BALB/c mice or ICR mice. Animals were then sacrificed after 0.25 h, 1 h, 2 h, 4 h, 8 h, 12 h and 24 h. For every sampling time point, five mice were sacrificed. Overall, 45 neutropenic mice were used. Phage concentration was once again measured in whole blood and spleen homogenates using a plaque assay and qPCR.

As with the BALB/c mice, we once again observed a notable decrease in active phage particles in blood from ICR mice **(Fig. 4C)**. The C_max_ of phages in ICR mice is 1.97 x 10^7^ PFU/mL, whereas the C_24_ is 280 PFU/mL. The *t*_1/2_ of phages in the blood of ICR mice is 3.66 h. In contrast, the concentration of active viral particles only decreased slightly in spleen tissue over the measured duration. The C_max_ of LPS-5 in the spleens of ICR mice was 3.89 x 10^8^ PFU/mL, whereas the C_24_ of phages in ICR mice was 5.31 x 10^6^ PFU/mL **(Fig. 4D)**. Other pharmacokinetic parameters are listed in **Table 1**.

As with the BALB/c mice, these data indicate that in neutropenic ICR mice, LPS-5 bacteriophages leave the circulation rapidly after administration and accumulate in other compartments like the spleen.

In comparing the figures for neutropenic ICR mice to those from conventional BALB/c mice we are struck that the figures for C_max,_ C_24,_ and *t*_1/2_ are very similar between these strains. For example, the *t*_1/2_ of phages in blood in BALB/c and ICR mice is 3.45 and 3.66 h, respectively. These data suggest that neutrophil-mediated phagocytosis is not a major determinant of phage clearance.

### Phages are inactivated in circulation

Our data reveal a substantial discrepancy in phage quantity over time when measured by qPCR versus plaque assay in blood. The C_24_ in the blood of the BALB/c mice measured by qPCR is 2.73 x 10^4^ PFU/mL versus 740 PFU/mL measured by plaque assay. This is a difference of around 30-fold. In the ICR mice, the C_24_ is 3610 PFU/mL measured by qPCR versus 280 measured by plaque assay. This is equivalent to a 12-fold difference, approximately (**Table 1**).

This observation is also supported by the disparity in the calculated *t*_1/2_. The *t*_1/2_ in the blood of the BALB/c mice measured by qPCR is 16.26 h versus 3.45 h measured by plaque assay. In the ICR mice, the *t*_1/2_ is 9.15 h measured by qPCR versus 3.66 h measured by plaque assay (**Table 1**).

These data indicate that factors in circulation reduce the number of functional phages (measured by plaque assay) compared to the total number of phages (measured by qPCR). The fact that the phages stay active for longer in the spleen than in the blood suggests that soluble factors in the blood inactivate phage particles. Additionally, we can conclude that phages can retain activity in spleen tissue over the measured time whereas their activity drops in the blood compartment.

## Discussion

Here, we have examined the role of neutrophils in bacteriophage pharmacokinetics using well-established mouse models and a phage from a recent human clinical trial. We observe that the clearance of phages from blood and spleen is only minimally affected by neutropenia. For example, the *t*_1/2_ of phages in blood in BALB/c and ICR mice is 3.45 and 3.66 h, respectively. These data suggest that phagocytosis is unlikely to be a major cause of phage clearance in the body.

These data are consistent with other literature suggesting that phagocytosis is not a major factor in phage clearance. Indeed, the small size of phage particles (25-250 nm) may be too small for conventional phagocytic mechanisms, as this typically involves particles of 2–3 μm^32^.

Our data do implicate serum factors in phage neutralization. We observe a substantial discrepancy in phage levels over time when measured by qPCR versus plaque assay. For example, the blood *t*_1/2_ of phages is 16.26 vs. 3.45 h when measured by qPCR or plaque assay, respectively. This suggests that substantial functional inactivation of circulating phages occurs over time, perhaps due to complement, fibrinogen, or other factors. To this point, there are reports indicating that bacteriophages are susceptible to complement-mediated inactivation via direct destruction or opsonization^33,34^. Since these mice had not seen LPS-5 phage before, it is extremely unlikely that they possessed neutralizing antibodies against these phages.

These studies have several implications for phage therapy. These data suggest that clearance from peripheral blood is fairly rapid and that local delivery may prove beneficial. They suggests that approaches to identify and nullify factors involved in phage clearance in circulation might promote better pharmacokinetics.

This work has several limitations. One limitation is that these studies do not include bacterial infection. The presence of infection and inflammation could dramatically impact the biodistribution and pharmacokinetics of phages. Moreover, since the phage population replicates itself in the presence of host bacteria, this element of increase through self-replication in phage populations is missing from our study. These issues and the impact of phage replication on pharmacokinetics were addressed in an excellent recent review^35^. Future studies will calculate the PK of phages in other tissue compartments beyond the spleen and blood and will extend this work to other phages and more complex dosing regimens.

## Acknowledgments

We gratefully acknowledge the following funding: National Institutes of Health grant R01 HL148184-01, R01 AI12492093, R01 DC019965, and grants from the Cystic Fibrosis Foundation (CFF), the Cystic Fibrosis Research Institute (CFRI) and the Emerson Collective grant (to PLB) and grants from the Stanford University Medical Scientist Training Program grant T32-GM007365 and the Stanford Interdisciplinary Graduate Fellowship, Gold Family Graduate Fellow (to TD) and the Merrigan fellowship (to AE).

## Author Contributions

Conceptualization: AE, PLB, Methodology: TD, AE, Investigation: TD, AE, and Writing: TD, AE, PLB

## Competing Interest Statement

RM is a member of Felix Biotechnology, Inc., a phage therapy company. All other authors declare they have no competing interests.

## Abbreviations

(*t*_1/2_): Half-life
(C_24h_): Initial concentration 0.25h after dose (C_max_) concentration at 24h after dose
(AUC_last_): area under the curve from time 0 to last measurable concentration
(AUC_Inf_): area under the curve from time 0 to infinity
(MRT): Mean residence time
(V_ss_): Volume of distribution at steady state
(CL): clearance

